# The *Escherichia coli* TolC efflux pump protein is immunogenic and elicits protective antibodies

**DOI:** 10.1101/2023.10.09.559948

**Authors:** TO Silva, LVS Costa, L Chagas do Nascimento, LS Barbosa, JGC Fanticelli, C Rotilho, LS Silva, J dos Santos Abreu, BF Rajsfus, ACS Bulla, J Carneiro, D Allonso, DR Salgado, ML da Silva, J Echevarria-Lima, LO Moreira, PC Olsen

**Author notes:** **Corresponding Authors: Priscilla Christina Olsen**, Laboratório de Estudos em Imunologia, Departamento de Análises Clínicas e Toxicológicas, Faculdade de Farmácia, Universidade Federal do Rio de Janeiro (UFRJ). Av. Carlos Chagas Filho 373, Bloco A 2° Andar sala 05, Cidade Universitária, Ilha do Fundão, CEP. 21941-902, Rio de Janeiro, Brazil.; **Lilian Oliveira Moreira**, Laboratório de Bacteriologia e Imunologia Clínica, Rua Professor Paulo Rocco, Bloco A2-07, Centro de Ciências da Saúde (CCS), UFRJ, Cidade Universitária, CEP: 21941-902, Rio de Janeiro, RJ, Brazil.

## Abstract

Antimicrobial resistance is an increasing worldwide public health burden that threatens to make the existent antimicrobials obsolete. Among the mechanisms of antimicrobial resistance is the overexpression of efflux pumps, such as the AcrA–AcrB–TolC which extrude diverse compounds, reducing the intracellular concentration of antimicrobials. TolC is the outer membrane protein of this pump and has recently gained attention as a therapeutic target. However, little is known about the immune response generated against the TolC protein. Here we evaluated the cellular and humoral immune response against the TolC from *Escherichia coli*. An *in silico* epitope prediction of the *E. coli* TolC showed several residues could bind to human antibodies, and we showed that human plasma presented anti-TolC IgG and IgA antibodies. Gram-negative infected patients presented a slight increase in anti-TolC IgM amounts, compared to controls. Recombinant *E. coli* TolC protein stimulated macrophages *in vitro* to produce nitric oxide, as well as IL-6 and TNF-α, assessed by Griess assay and ELISA, respectively. Immunization of mice with TolC intraperitoneally and an *in vitro* re-stimulation of lymph node cells led to increased percentage of T cell proliferation and IFNγ production, evaluated by flow cytometry and ELISA, respectively. We observed that TolC mouse immunization stimulated anti-TolC IgM and IgG production, with a higher level of IgG1 and IgG2, amongst the IgG subclasses. Finally, TolC IgG from mouse immune serum could bind to live *E. coli,* increase bacterial uptake by macrophages *in vitro*. TolC immunized mice had a survival rate increased in 60% post infection with *E. coli*. Our results showed that TolC is immunogenic, activating macrophages, T and B cells, leading to the production of protective antibodies against *E. coli*.

## Introduction

Antimicrobial resistance occurs when bacteria display the ability to bypass antimicrobial pharmacological mechanisms used against them and is mainly associated with the widespread use or misuse of antimicrobials in human therapy, and also in agriculture, livestock and industry. Infections caused by antimicrobial-resistant pathogens are more difficult to treat and can cause significant morbidity and mortality ^1,2^. It is estimated that, globally, 700,000 deaths annually are attributed to antimicrobial resistant microorganisms and this number could increase to 10 million deaths per year by 2050, surpassing current leading causes of death, including cancer ^3^. In 2017, the World Health Organization (WHO) provided a priority list of antimicrobial-resistant bacteria to guide research and development of new antimicrobials against them^4^. *Escherichia coli* (*E. coli*) was one of the Gram-negative bacteria ranked as a critical priority in this context*. E. coli* is a gut commensal bacterium, however, several pathotypes can be isolated from a wide range of infections in humans and animals, such as urinary tract infections, meningitis, bacteremia and bowel disorders, ranging from mild diarrhea to severe colitis. In general, antimicrobial resistance in *E. coli* is considered one of the greatest challenges in humans and veterinary medicine worldwide, with great economic burden, and needs to be considered a public health concern, especially in a One Health perspective^5^.

Antimicrobial resistance is typically associated to bacteria ability to degrade or modify of the antibiotic, change the drug target, and reduce antimicrobial accumulation within the cell due to decrease of drug permeability or increase of efflux pumps production^6^. Efflux pumps are specialized protein complexes found in the membranes of bacterial, fungal and mammalian cells, that function to remove certain substances from the cell by actively pumping them out of the cell. In bacteria, efflux pumps play a significant role in antimicrobial resistance, as they can pump out antimicrobials before the drugs reach their target site and kill the bacteria ^7^. It has already been shown that the presence of antimicrobials can induce overexpression of efflux pumps in the bacterial membrane, leading to bacterial resistance ^8,9^. Interestingly, since this mechanism decreases the intracellular concentration of antimicrobials, it may contribute to the development of other resistance mechanisms leading to multidrug-resistance (MDR) ^10,11^.

Five different families of efflux pumps were identified in *E. coli*, the ATP-binding cassette (ABC) superfamily, major facilitator superfamily (MFS), multidrug and toxic compound extrusion (MATE) family, small multidrug resistance (SMR) family and resistance-nodulation-cell division (RND) family. The AcrA–AcrB–TolC RND-tripartite complex constitutes a key efflux pump related to bacterial resistance, able to extrude a wide range of antimicrobials as well as toxic chemicals, such as a variety of detergents and dyes ^12–14^. The protein TolC can interact with inner membrane components of RND, ABC and MFS efflux pumps. TolC assembles as a trimer in the outer membrane, forming a pore, and presents two extracellular residues ^15,16^. *E. coli* bacterial growth was inhibited by serum from TolC-immunized rabbits in the presence of 1/8 MIC antibiotic treatment and TolC from *Edwardsiella tarda* whole-cell vaccine was found to be an important immunogen with protective effects ^17,18^. Therefore, TolC can be a potential therapeutic target and vaccine candidate mainly against Gram-negative resistant bacteria. However, little is known about the immune response against the TolC efflux pump protein in mice and humans. In this work we aim to characterize the cellular and humoral immune response against the *E. coli* TolC protein and find antibodies that present protective effects.

## Methods

### TolC cloning, expression and purification

The *E. coli* (strain K12) TolC gene (Uniprot P02930) was synthesized by GenScript (USA) and inserted into a pET30a vector. A His-Tag and GFP-tag were inserted downstream TolC gene before the stop codon. *E. coli* BL21(λDE3) cells transformed with pET30-TolC were grown in Luria Bertani (LB) broth with 50 µg/mL kanamycin at 37°C until reaching an optical density at 600 nm (OD_600nm_) of 0.6. TolC recombinant expression was induced with 1 mM of isopropyl β-D-1-tiogalactopiranosideo (IPTG) followed by cultivation overnight at 15°C. Cell culture was then centrifuged at 5000 g for 15 min at 4°C and the pellet was resuspended in 50 mM Tris-HCl pH 8, 50 mM NaCl, 1 mM DTT, 0.1% CHAPS, and 5% glycerol with protease inhibitors cocktail (Sigma Aldrich, USA). The cells were lysed with 5 mg/mL of lysozyme and chromosomal DNA was cleaved by addition of 20 mg/mL of DNAse and 5 mM MgCl_2_. The suspension was then sonicated at 40% amplitude for 10 cycles (15 s on/45 s off), followed by centrifugation for 20 min at 20000 g at 4°C.

Inclusion bodies containing recombinant TolC were resuspended in 50 mM Tris-HCl pH 8, 50 mM NaCl, 1 mM DTT, 0.1% CHAPS, 5% glycerol and 8 M urea overnight at 4°C with constant stirring, followed by centrifugation at 20000 g at 4°C. Supernatant was collected and purified in a Ni^+2^NTA affinity column (HisTrap HP– GE Healthcare, USA) according to manufacturer protocol. The purified protein was dialyzed in 50 mM Tris-HCl pH8.0, 50 mM NaCl, 1 mM DTT, 0.1% CHAPS, and 10% glycerol and stored at -80°C.

Protein purity was about 90% (densitometric analysis of Coomassie Blue-stained SDS-PAGE gel under reducing condition) and LPS content was evaluated with Pierce^TM^ Chromogenic Endotoxin Quant Kit (Thermo) (data not shown).

### Epitope prediction from *E. coli* TolC

The protein sequence of *E. coli* TolC was retrieved from UniProtKB and the protein external region was assessed for antigenicity using the Vaxijen 2.0 web server ^19^, with the target organism set to “bacteria” and a threshold of 0.5. As flexible regions in the protein are often associated with antigenicity^20^, the Karplus & Schulz Flexibility Prediction tool from IEDB was employed to predict flexibility. Furthermore, the IUPred3 tool predicted intrinsically disordered regions^21^.

Multiple tools were used to predict the location of linear B-cell epitopes: (i) BepiPred-2.0 on the IEDB server, employed with a default threshold of 0.5, using a Random Forest algorithm trained on epitopes and non-epitope amino acids from crystal structures to predict the location of linear B-cell epitopes ^22^, (ii) ABCPred tool^23^ was utilized with a threshold of 0.21, which is a neural network specifically trained to identify linear B-cell epitopes, Nad (iii) the Epitopia algorithm^24^ was used, which is a machine-learning tool trained to identify antigenic features. To predict conformational B-cell epitopes, the crystallographic structure coding PDBid 1TQQ from the Protein Data Bank – PDB was used. The protein structure was refined using GalaxyRefine^25^, which involves reconstructing the side chains and relaxing the structure through molecular dynamics simulations. The stereochemical analysis of the PDBid 1TQQ was conducted using MolProbity^26^. For conformational epitope prediction, DiscoTope^27^ and Ellipro^28^ were used, with threshold values of -3.7 and 0.5, respectively. Additionally, we employed the Epitopia tool for this analysis^24^.

To assess the binding affinity of peptides to multiple human leukocyte antigen (HLA) class II molecules, the Major histocompatibility complex class II (MHC II) binding prediction tools of IEDB server NN-align2.3 (netMHCII-2.3) were used. NN-align, an artificial neural network-based alignment method was used to calculate the IC_50_ values for peptides binding to specific MHC molecules, where low IC_50_ values indicate good binders^29^. The NN-align method was developed using a quantitative binding data set of MHC class II peptides in IEDB databases. The predicted length of CD4+ T cell epitopes bound to MHC class II molecules was set to 15 amino acids. The set of alleles (DRB1*03:01, DRB1*07:01, DRB1*15:01, DRB3*01:01, DRB3*02:02, DRB4*01:01, and DRB5*01:01), was selected since they best represent the human population for epitope prediction tools^30^.

### Macrophage stimulation with TolC *in vitro*

RAW 264.7 murine macrophage lineage (0.5 x 10^5^/well in 24 well plate) was cultured in DMEM (Gibco/Thermo) supplemented with 10% inactivated FCS (Gibco/Thermo), 1% penicillin/streptomycin (Gibco/Thermo) and NaHCO_3_ in the absence or presence of GFP^+^ recombinant TolC (1, 10 and 100 μg/mL) for up to 72h at 37 °C and 5 % CO_2_. LPS (0.5 μg/mL) from *E. coli* serotype 0111:B4 (Sigma-Aldrich/Merck) was used as a control of macrophage stimulation. GFP fluorescence on macrophages was detected at 3 and 24h post stimulation through flow cytometry (FACS Calibur; BD Biosciences). Supernatants were collected at 24 and 72h for evaluation of cytokines and nitric oxide (NO) production, respectively. NO production was determined by the detection of nitrite using the Griess method adapted ^31^. On assays including polymyxin B (1 μg/mL) incubated with either 10 ng/mL LPS or 10 μg/mL of TolC, supernatant was collected at 72h for evaluation of NO detection.

### Mouse experimental assays

Eight-week-old (18–20g) female or male BALB/c and C57BL/6 mice were obtained from the FECD/Biotério Decania UFRJ Facility (Rio de Janeiro, Brazil). Mice were kept in the animal-housing facilities at a controlled room temperature (22–25 °C) and a 12-h (6 am–6 pm) light– dark cycle. Procedures involving the care and use of laboratory animals were examined and approved by the Animal Ethics Committee of the Federal University of Rio de Janeiro (CEUA A04-21-109-19 and 017-22). Weight and survival were monitored over time in all procedures. Serum of germ-free mice were kindly donated by Leda Quercia Vieira and Flaviano dos Santos Martins (UFMG).

BALB/c or C57/BL6 mice were immunized on day 0 by an intraperitoneal injection (0.2 mL) of a mixture containing TolC recombinant protein (10 μg) and Imject^TM^ Alum in 0.9% NaCl sterile solution (saline). Booster was performed in BALB/c mice 14 days after the first immunization, to retrieve serum and memory B cells at 28 (days post immunization) dpi. Blood samples were collected on 0, 7, 14 and 35 dpi. A group of mice were culled (100 mg/kg Thiopental plus 10 mg/mL lidocaine on days 0, 7, 14 and 35 to collect total lymph nodes and spleen. In another set of experiments mice were sensitized with the whole heat inactivated *E. coli* 25922 at days 0, 15 and 29, either intranasally (50 μL with 1 x 10^8^ CFU), intraperitoneally (200 μL with 1 x 10^8^ CFU) and orally (100 μL of 1 x 10^8^ CFU) diluted in sterile PBS. Blood and bronchoalveolar lavage (BAL) samples were retrieved at 36 dpi. On the assays performed to test the effect of immunization against infection, C57/BL6 mice were sensitized and boosted with TolC, as previously described, and 14 after the booster, mice were infected with 2 x LD_50_ of *E. coli* 25922 (6 x 10^7^ CFU/mouse ip) ^32^.

### Detection of TolC^+^ memory B cells

B cell labeling was performed as previously described in detail ^33^, with the following modifications. BALB/c mouse lymph nodes (cervical, inguinal, brachial and axillary) and spleen were retrieved and single-cell suspension was obtained through a 40 μm cell strainer, washed with PBS 3% FBS, and incubated at 4°C for 20 min with GFP-labeled TolC at 680 μg/mL and the following monoclonal antibodies: PE B220 at 1/200 (Thermo), PerCP eFluor 710 IgM at 1/200 (Thermo), BV421 IgG1 at 1/25 (BD Horizon^TM^), APC CD38 at 1/300 (Thermo), Dump channel (APC EF780 CD4, CD8, CD11c, CD11b, F4/80, Gr1 at 1/200, all from Thermo) and. After the last wash, TolC^+^ memory B cells were analyzed using a FACSAriaII (Becton Dickinson).

### Lymphocyte stimulation *in vitro*

The proliferation of lymph node (cervical, brachial, axillary and inguinal) cells from TolC or non-immunized mice was evaluated on 0, 14 and 35 dpi. Cells were collected from the lymph nodes, challenged with immobilized anti-CD3 (10 μg/mL) or soluble TolC (1, 10 or 100 μg/mL) *in vitro* for 72h and permeabilized and stained cells with propidium iodide (PI) as described previously ^34^. Briefly, cells were stained with PI (75 μM) in the presence of NP-40. Analyses of the DNA content were carried out by collecting 10,000 events for cell-cycle analysis using a FACSCalibur flow cytometer and FlowJo (BD Biosciences). Proliferation values were determined by calculating the percentage of cells in phase S+G2 of the cell cycle. Supernatant was also collected at 72h for evaluation of cytokine production.

### Collection of human sera

Human serum was obtained from samples of peripheral blood collected from Hospital Clementino Fraga Filho/UFRJ participants (CAAE: 48917121.0.0000.5257), upon informed consent. Participants from the ICU were included in the study after presenting Gram-negative bacteria in samples such as blood, urine or tracheal secretion. Individuals that did not present Gram-negative infection were used as controls. Serum aliquots were heat-inactivated at 56 °C for 1 hour and stored at -20 °C thereafter. The control group (CO) participants had an average age of 55.41 years old and 25% were male. The patient group (PT) was constituted of participants with the average age of 58.17 years old and 33.33% were male.

### Detection of mouse serum IgG binding to live bacteria

To determine the serum antibody binding to live bacteria, *E. coli* ATCC 25922 (3.75 x 10^6^) were stained with Syto9 500 nM (Invitrogen/Thermo) and incubated with non-immune or TolC-immune serum (1:100) for 15 min at 37 °C. Bacterial cells were washed and incubated with BV421 rat anti-mouse IgG1 at 1:25, (BD Horizon^TM^) for 30 min at 4 °C. Finally, the bacteria cells were fixated with 1% paraformaldehyde, and the fluorescence was detected by BD LSRFortessa^TM^ FACS and analyzed with FlowJo.

### TolC detection by Western blot using mouse immune serum

To obtain the bacterial cell proteins extract, *E. coli* ATCC 25922*, Acinetobacter baumannii* ATCC 19606*, Klebsiella quasipneumoniae* ATCC 700603 and *Pseudomonas aeruginosa* ATCC 27853. Bacteria were grown in Tryptic Soy Broth (TSB) medium, suspended in 4 mL of PBS (OD_600_ = 1.0), centrifuged (13.000g for 10 min), pellets mixed with lysis buffer (50 mM Hepes pH 7.5, 300 mM NaCl, 1 mM DTT, 0.01% CHAPS, 5% glycerol), and then boiled with sample buffer (Tris-Cl/SDS pH 6.8, 30% glycerol, 0.1% SDS, 0.09% DTT, 1.2% bromophenol blue) for 10 min at 96 °C. Total bacterial protein extracts (20 μL) or purified recombinant TolC protein (1 mg) electrophoretically separated by SDS-PAGE, transferred to nitrocellulose membranes that were blocked with Tris-buffered Tween 20 (TBS-T) containing 3% dry milk overnight at 4 °C. Membranes were incubated with mice TolC-immune or non-immune serum (1:1000) for 1 h at room temperature. Then, a rabbit polyclonal anti-mouse IgG (1:10.000) (Jackson Immunoresearch) was added for 1 h at room temperature. Finally, the positive bands were detected by chemiluminescence (ECL, Pierce) using a ChemiDoc^TM^ Imaging Systems^5,6^.

### *E. coli* macrophage uptake

The macrophage uptake test was performed to investigate the opsonization role of TolC immune serum on *E. coli*. RAW 264.7 macrophages were cultured in DMEM, as described previously. *E. coli* ATCC 25922 (3.75 x 10^6^ CFU/well) were stained with Syto9 500 nM and incubated with non-immune or TolC-immune serum (1:20) for 15 min at 4 °C. Bacterial cells were washed and incubated with RAW macrophages (7.5 x10^4^/well) for 30 min at 37 °C in DMEM media without FCS or penicillin/streptomycin. RAW cells were then washed twice, trypsinized, acquired in the flow cytometer BD LSRFortessa^TM^ and analyzed with FlowJo.

### ELISA assays

#### Anti-TolC ELISA

The binding of mouse or human serum (or BAL, where indicated) IgG, IgG subclasses, IgA and IgM antibodies to TolC recombinant proteins was assayed by ELISA. Plates were coated with 380 ng of TolC protein in PBS per well and incubated overnight at room temperature. Plates were then blocked with 1% BSA, 0.1 mM EDTA in PBS-T (PBS with 0.05% Tween20) for 1 h at 37 °C and washed with PBS-T in between each step. Serum samples were diluted with PBS-T and added for 1 h at 37 °C. For mouse samples, the secondary HRP-conjugated goat anti-mouse IgG, IgM or IgG2c (Jackson Immunoresearch), IgG1 or IgG3 (Thermo), IgG2b or IgA (Southern Biotech) (at the concentrations indicated by the manufacturer) were added for 1 h at room temperature. For human samples, the secondary HRP-conjugated goat anti-human IgG or IgA (Jackson Immunoresearch), IgG1, IgG2, IgG3, IgG4 or IgM (Southern Biotech) (at the concentrations indicated by the manufacturer) were added for 1 h at room temperature. Plates were then developed using ABTS substrate (Thermo) and read at 405 nm. The relative binding affinity of antibodies was determined similarly, using serially diluted samples or by calculating the ratio over negative control samples.

#### Anti-LPS ELISA

The binding of mouse or human serum IgG and IgM antibodies to LPS from *E. coli* serotype 0111:B4 (Invivogen) was evaluated by ELISA. Plates were coated with 10 μg/mL LPS, stored overnight at room temperature, and blocked with 1% BSA, 0.5 mM EDTA in PBS-T (PBS with 0.05% Tween20) for 2 h at room temperature. Plates were washed with PBS-T in between each step. Human or mouse serum samples were diluted at 1:640 or 1:40, 1:200 and 1:1000 with PBS-T and added for 2 h at room temperature. Secondary goat HRP-conjugated anti-mouse or anti-human were added, following 1h incubation at room temperature. Plates were developed as described above.

#### Detection of cytokines

Secretion of IL-6 and TNF-α was detected in the supernatant of RAW macrophage culture upon stimulation with either LPS or TolC as described previously. Secretion of IFN-γ was detected in the supernatant of the cell culture performed with cells obtained from total lymph nodes of non-immunized mice or TolC-immunized mice (14 and 35 days post immunization). Lymph node cells were re-stimulated *in vitro* with either anti-CD3 or TolC for 72h. Analyses of the production of cytokines were undertaken as indicated by the ELISA kit manufacturer (Thermo).

### Statistical analysis

Statistical analyses were performed with Prism 7.0 software, carried out with ordinary one way-ANOVA with Tukey’s multiple comparisons test or with unpaired t test in between two groups. P < 0.05 was considered significant. The two-tailed Pearson r test correlation was used for the correlations of figure S1. The values of n and what they represent, as well as symbols representing P value, are described in each figure legend.

## Results

### *In silico E. coli* TolC epitope prediction

The extracellular exposed regions of TolC protein “GYRDANGI” and “DTSYSGSKTRGAAGTQYDDSN” showed antigenicity predicted by VaxiJen of 0.8484 and 1.2808, respectively (Figure 1A and B). The flexibility score of TolC exceeded 1.0 and the intrinsic disorder were identified in the exposed regions, indicating the presence of potential antigenic determinants ^20^ and a propensity for protein-protein interactions ^21^. The linear prediction programs identified potential B cell epitopes in the exposed regions of the protein (Figure 1C). BepiPred-2.0 identified all residues as epitopes. In ABCPred with a score of 0.88, “DYTYSNGYRDANGINS” and 0.90, “AGTQYDDSNMGQNKVG” were identified as potential epitopes. In Ellipro, “LGLGADYTYSNGYRDANGINSNATSASLQL” with a score of 0.838 and “IRQAQDGHLPTLDLTASTGISDTSYSGSKTRGAAGTQYDDSNMGQNKVGLSFSLPIYQGGMVNSQVKQAQYNFVGASE” with a score of 0.753 were identified as potential epitopes (Figure 1C). The trimeric structure of TolC (PDBid 1TQQ) was refined with GalaxyRefine and evaluated with MolProbity. The Clashscore (1.98) and MolProbityScore (1.00) values indicated a low presence of stereochemical issues. The exposed residues were positive for an epitope in the analysis with Discotope, Ellipro, and Epitopia (Figure 1). The highest Propensity Score from Discotope was 3.939, and the lowest was -3.248, for a threshold of – 3.7. In Ellipro, the lowest score per residue was 0.886, and the highest was 0.998. With Epitopia the average score of the residues was 0.703 for region 1 and 0.686 for region 2.

**Figure 1:**
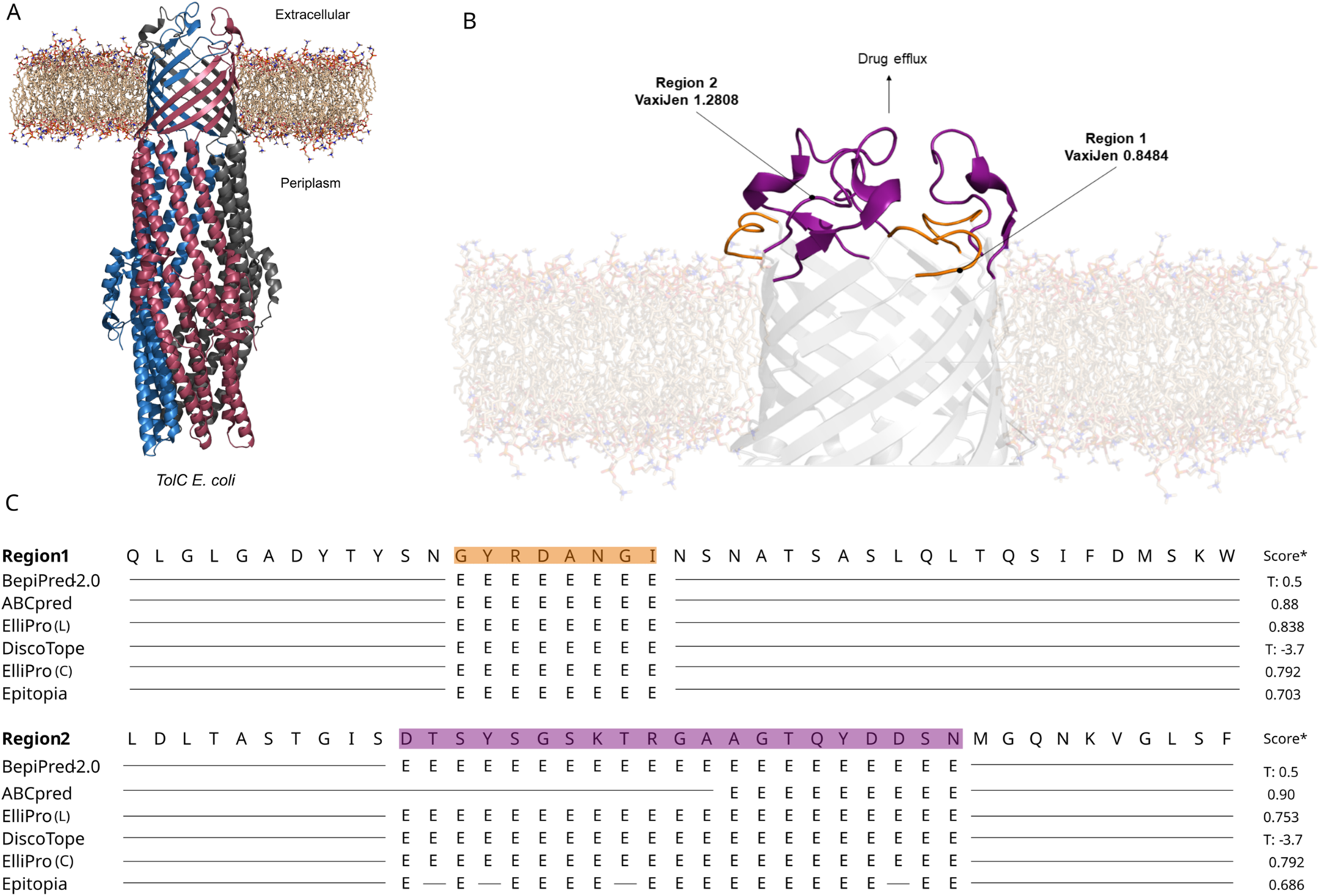
Structure, location, and immunogenicity prediction of *E. coli* TolC. (A) Representation from the homotrimer structure of outer membrane protein TolC from *Escherichia coli* (PDB id 1TQQ). Each color (gray, pink, and blue) is a different chain. (B) Region of TolC facing the extracellular environment. The antigenicity prediction with VaxiJen for chain A of the protein was 0.8484 (region 1) and 1.2908 (region 2). (C) B-cell linear and conformational epitope prediction. For region 1 (orange) and region 2 (purple), potential linear epitopes (BepiPred, ABCPred, and ElliPro) and conformational epitopes (Discotope, ElliPro, and Epitopia) were identified. The prediction results indicate a propensity for immune system recognition of these regions. ElliPro (L): linear epitope. ElliPro (C): conformational epitope. Score*: score assigned by each respective prediction program. T: Threshold. E: predicted epitopes.

For the identification of helper T-cell epitopes, the NN-align2.3 (netMHCII-2.3) was used, which predicts MHCII peptide binding through artificial neural networks. Helper T cells play a crucial role in adaptive immunity. Several sites along the protein generated high-affinity peptides for the most common MHC-II alleles, in the external region of TolC, the identified epitopes had intermediate (IC_50_ < 500 nM) and low affinity (IC_50_ < 5000 nM) classifications. For other regions of the protein, 115 epitopes were identified with high affinity (IC_50_ <50 nM) (Supplementary Table 1).

### Human plasma presents anti-TolC antibodies

Human plasma collected from either control individuals or from UCI Gram-negative infected patients were assayed to evaluate the presence of antibodies against TolC. All individuals presented antibodies against TolC, with higher IgG and IgA anti-TolC titers than IgM (Figure 2A). However, higher IgM anti-TolC titers were detected in patient plasma than in controls (Figure 2B), while anti-TolC IgG levels were not different in between these two groups (Figure 2C). Anti-TolC IgA plasmatic titers were also increased both in both control and patient groups, when compared to IgM (Figure 2A and 2D). The anti-TolC IgG subclasses of were also tested, and no difference was observed between the control and patient groups (Figure 2E, F, G and H). IgG2 was the predominant subclass of anti-TolC IgGs in humans (controls and patients).

**Figure 2:**
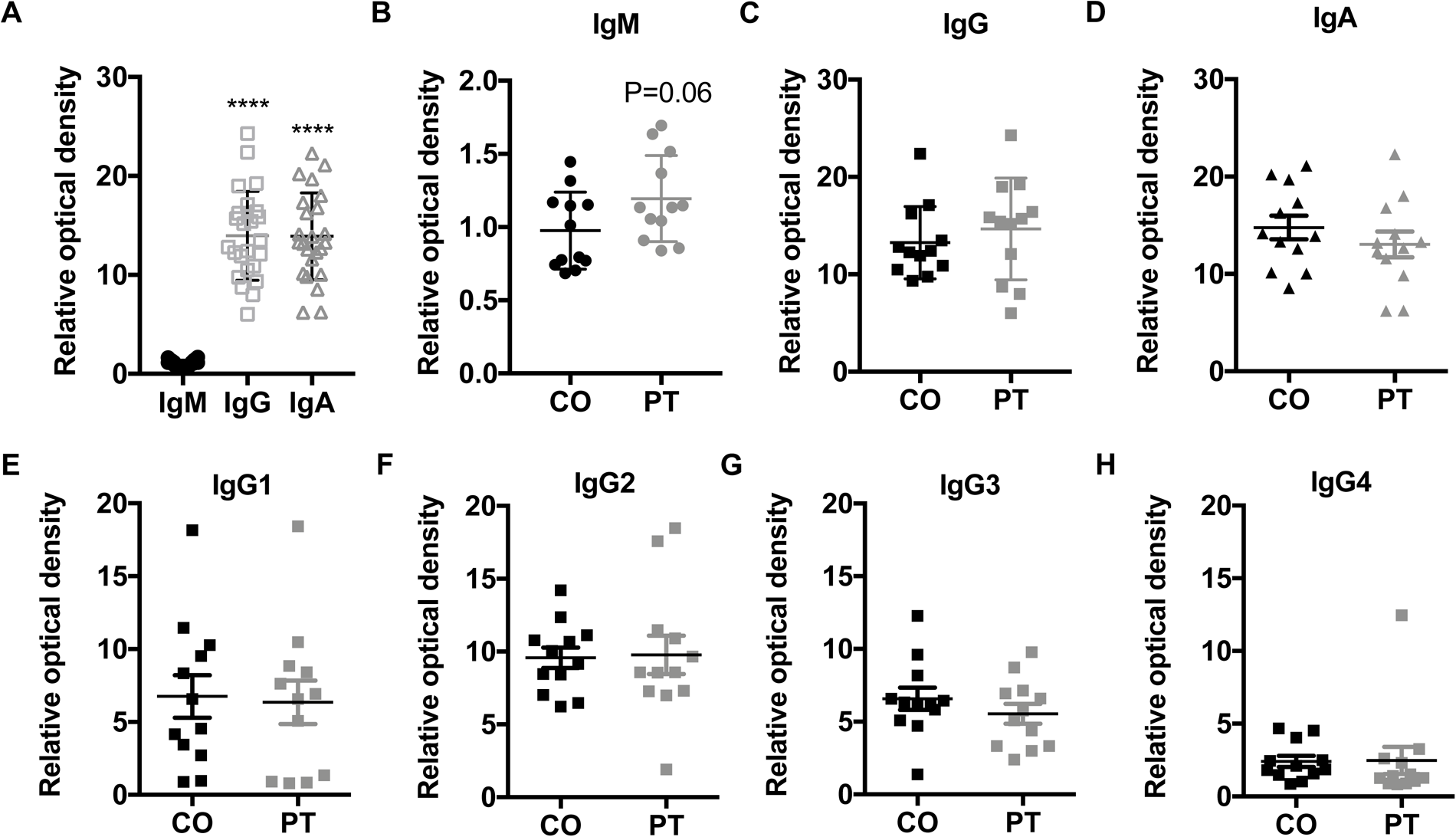
Anti-TolC antibodies are present in human plasma. (A) Anti-TolC IgG and IgA are more abundant than IgM in human plasma, when analyzed controls and patients together. (B) Anti-TolC IgM in controls or patients recently infected with Gram-negative bacteria. (C)Anti-TolC IgG or IgA (D) in human plasma from patients or controls. (E) Anti-TolC IgG1,(F) IgG2, (G) IgG3 andIgG4 (H) in human plasma from patients or controls. Control (CO) or patient (PT) groups N=12. **** P <0.0001compared to IgM.

Although anti-TolC IgM seemed to be increased in patients after/during a Gram-negative infection, there was no difference in anti-LPS IgM or IgG titers (Figure 1S A and B). Also, there was no significant difference in the amounts of anti-TolC IgM or IgG when results were segregated by gender (Figure 1S C and D). Anti-TolC IgM was not correlated with age in neither group analyzed, but IgG anti-TolC was negatively correlated with age in the patient group (Figure 1S E and F). IgG anti-TolC was correlated to anti-TolC IgM in the control group, but not in the patient group (Figure 1S G), possibly due to the anti-TolC IgM increase in the latter group (Figure 1S H). These results suggest TolC protein is immunogenic in humans and anti-TolC antibodies could be related to infection by Gram-negative bacteria.

### TolC recombinant protein activates macrophage *in vitro*

RAW macrophage lineage cells stimulated with 10 or 100 μg/mL GFP TolC *in vitro* presented an increase of GFP fluorescence at 3h of incubation, when compared to control cells (Figure 3A). At 24h of incubation, GFP fluorescence could not be detected anymore (data not shown). Macrophage stimulated with TolC *in vitro* (10 or 100 μg/mL) produced NO (Figure 3B) and the inflammatory cytokines IL-6 and TNF-α, (Figure 3C and D). Increase in IL-6 and TNF-α induced by TolC was concentration dependent. The production of NO, IL-6 and TNF-α induced by 100 μg/mL of TolC was comparable to what was observed after LPS stimulation (0.5 μg/mL). The effect of TolC on macrophage was not due to a possible LPS contamination, since LPS blockage with polymyxin B did not revert the TolC effect (Figure 3 E). These results demonstrate TolC recombinant protein is recognized by macrophage and leads to their activation.

**Figure 3:**
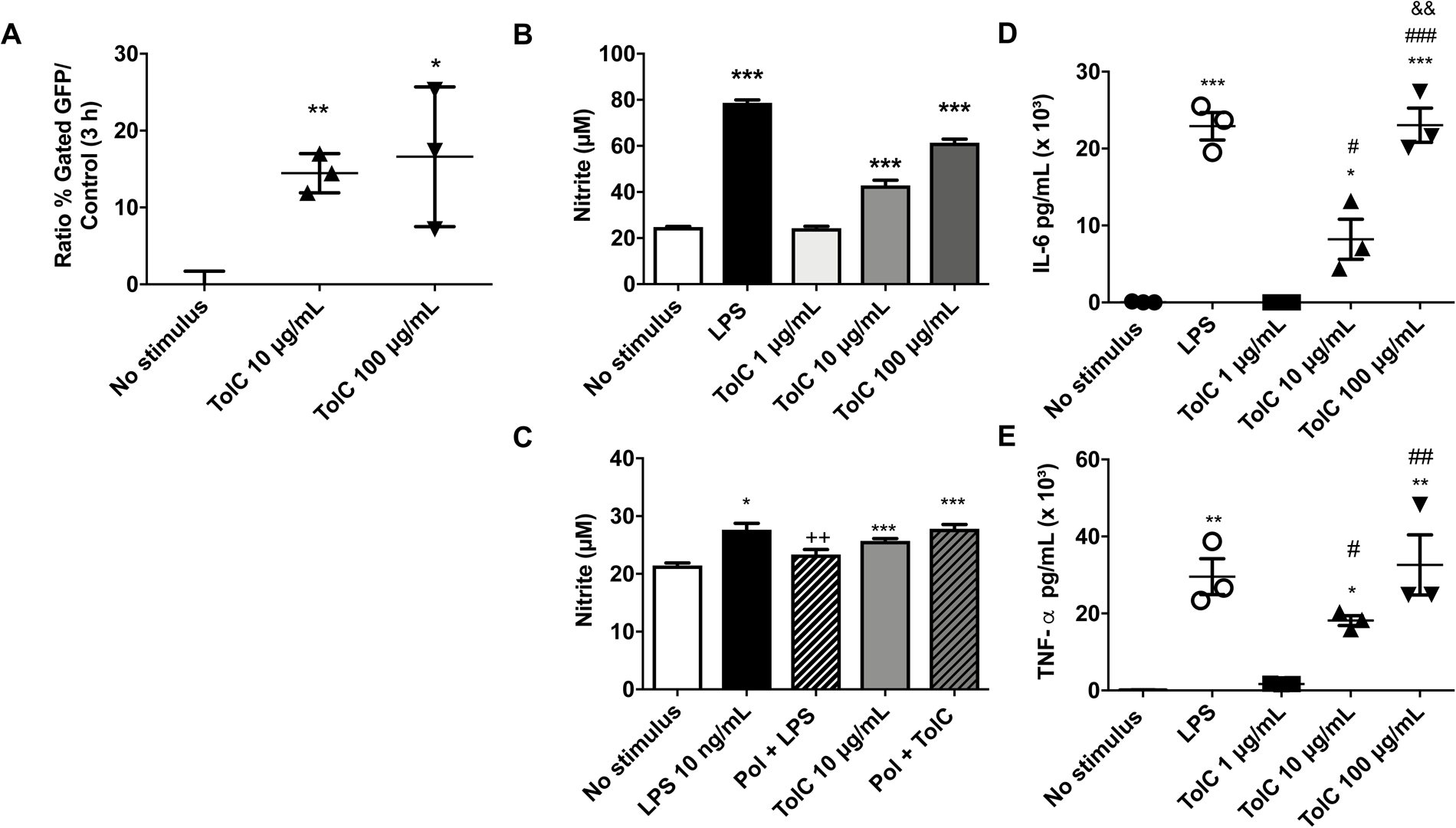
TolC recombinant protein induces macrophage activation *in vitro*. (A) Macrophage fluorescence detection 3 h after TolC-GFP stimulus, detected by flow cytometry. (B) Nitric oxide production on macrophage supernatant 72h after LPS (0.5 μg/mL) or TolC stimulation (1, 10 and 100 μg/mL). (C) Nitric oxide production on macrophage supernatant 72h after polymyxin B (1 μg/mL) plus LPS (10 ng/mL) or TolC stimulation (10 μg/mL). (D) IL-6 and (E) TNF-a secretion on macrophage supernatant 24h after LPS or TolC stimulation. * P <0.05, ** P <0.01 and *** P <0.001 compared to cells with no stimulus. ^#^ P <0.05, ^##^ P <0.01 and ^###^P <0.001 compared to cells stimulated with 1 μg/mL of TolC. ^&&^P <0.01 compared to cells stimulated with 10 μg/mL of TolC.

### Mouse immunization with TolC leads to lymphocyte activation

Next, we sought to test if TolC recombinant protein was able to activate lymphocytes. To investigate that, C57/Black6 mice were immunized with TolC intraperitoneally and their total lymph nodes were retrieved at 14 and 35 dpi in order to evaluate *in vitro* lymphocyte proliferation in the presence of antigenic recall with TolC. Non-immunized mice did not present lymphocyte proliferation *in vitro* when stimulated with 1, 10 or 100 μg/mL of TolC, although there was significant proliferation induced by anti-CD3, when compared to non-stimulated cells (Figure 4A). At 14 dpi cells from immunized mice re-stimulated with TolC (100 μg/mL) presented increased proliferation, as observed in anti-CD3 stimulated cells (Figure 4B). After 35 dpi the proliferation induced by TolC (100 μg/mL) *in vitro* was still present, but it was diminished when compared to 14 dpi (Figure 4C).

**Figure 4:**
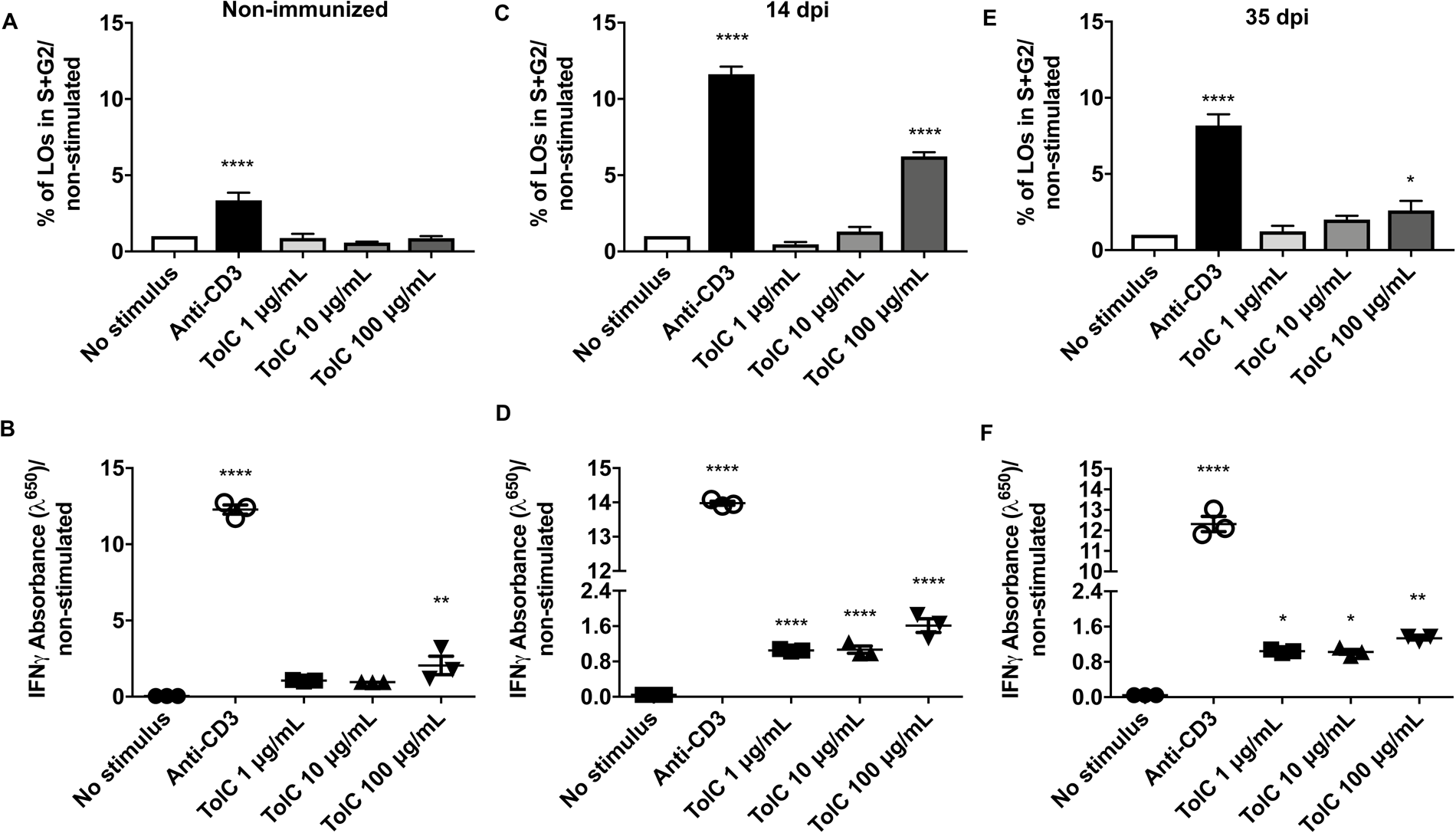
Immunization of mice with TolC recombinant protein induces lymphocyte activation upon *in vitro* antigenic recall. Lymph node cells from non-immunized mice proliferation (A) and IFN-γ secretion (B) 72h after anti-CD3 or TolC *in vitro* stimulation (1, 10 and 100 μg/mL). Lymph node cells from TolC immunized mice (14 dpi) proliferation (C) and IFN-γ (D) secretion 72h after anti-CD3 or TolC *in vitro* stimulation. Lymph node cells from TolC-immunized mice (35 dpi) proliferation (E) and IFN-γ (F) secretion 72h after anti-CD3 or TolC*in vitro* stimulation. * P <0.05, ** P <0.01 and **** P <0.0001 compared to cells with no stimulus *in vitro*. C57/BL6 mice N=3.

Lymph node cells from non-immunized mice stimulated *in vitro* with anti-CD3 or TolC (100 μg/mL) presented increased secretion of IFN-γ, compared to non-stimulated cells (Figure 4D). Lymph node cells obtained at 14 and 35 dpi re-stimulated with anti-CD3 or TolC at all concentrations presented increased IFN-γ secretion, compared to non-stimulated cells (Figure 4 E and F). At all-time points evaluated, TolC did not induced IL-10 secretion (data not shown). These results demonstrate that TolC recombinant protein is able to stimulate lymphocytes leading to increased proliferation and pro-inflammatory cytokine production.

### Immunization with TolC protein or whole cell *E. coli* increases anti-TolC antibody production

Immunization of mice with TolC led to an increase in anti-TolC IgM at 7 dpi, which decreased by 10 dpi (Figure 5A). Immunized mice presented an increase of anti-TolC IgG titers on the 10^th^ dpi that was still high at 35 dpi, when compared to non-immunized mice (Figure 5B). Anti-TolC IgM titers were increased in serum from immunized mice at 7 dpi, when compared to non-immunized mice (Figure 5A and C). Interestingly, at 1:20 serum dilution there was anti-TolC IgM detection in non-immunized mice, which was not observed in germ-free mice (Figure 5C). At 14 dpi immunized mice presented higher anti-TolC IgG titers than non-immunized and Germ-free mice (Figure 5D). Non-immunized mice presented more IgG anti-TolC at 1:20 serum dilution when compared to Germ-free mice (Figure 5D). Anti-TolC IgG subclasses IgG1, IgG2b and IgG2c were produced after 15 dpi, but not IgG3 (Figure 5 E, F, G and H). TolC intraperitoneal immunization did not elicit serum anti-TolC IgA (Figure 5 I) or lead to increased anti-LPS IgM or IgG (Figure S2). There was also no significant loss of weight or survival impact due to the immunization (Figure S3 A and B).

**Figure 5:**
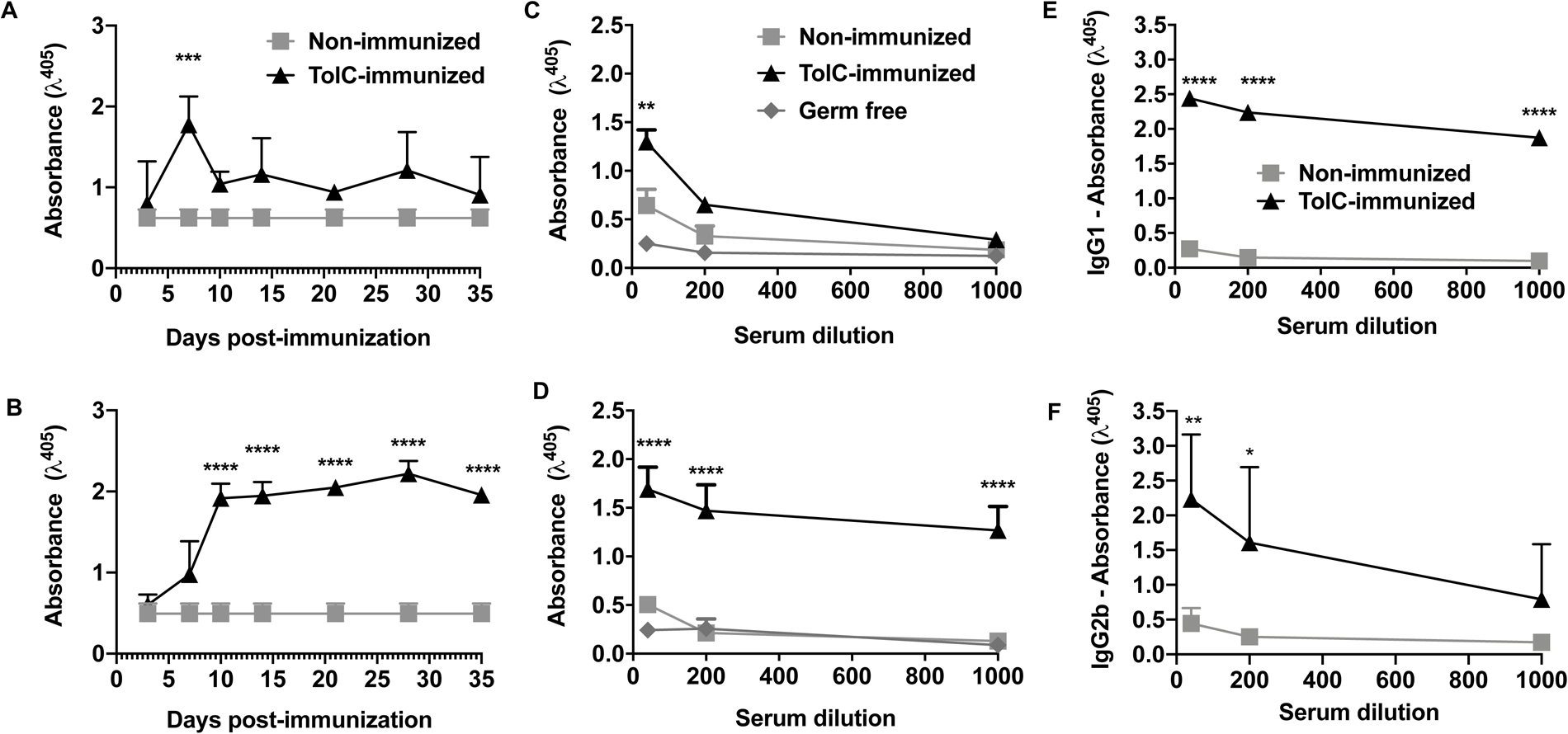
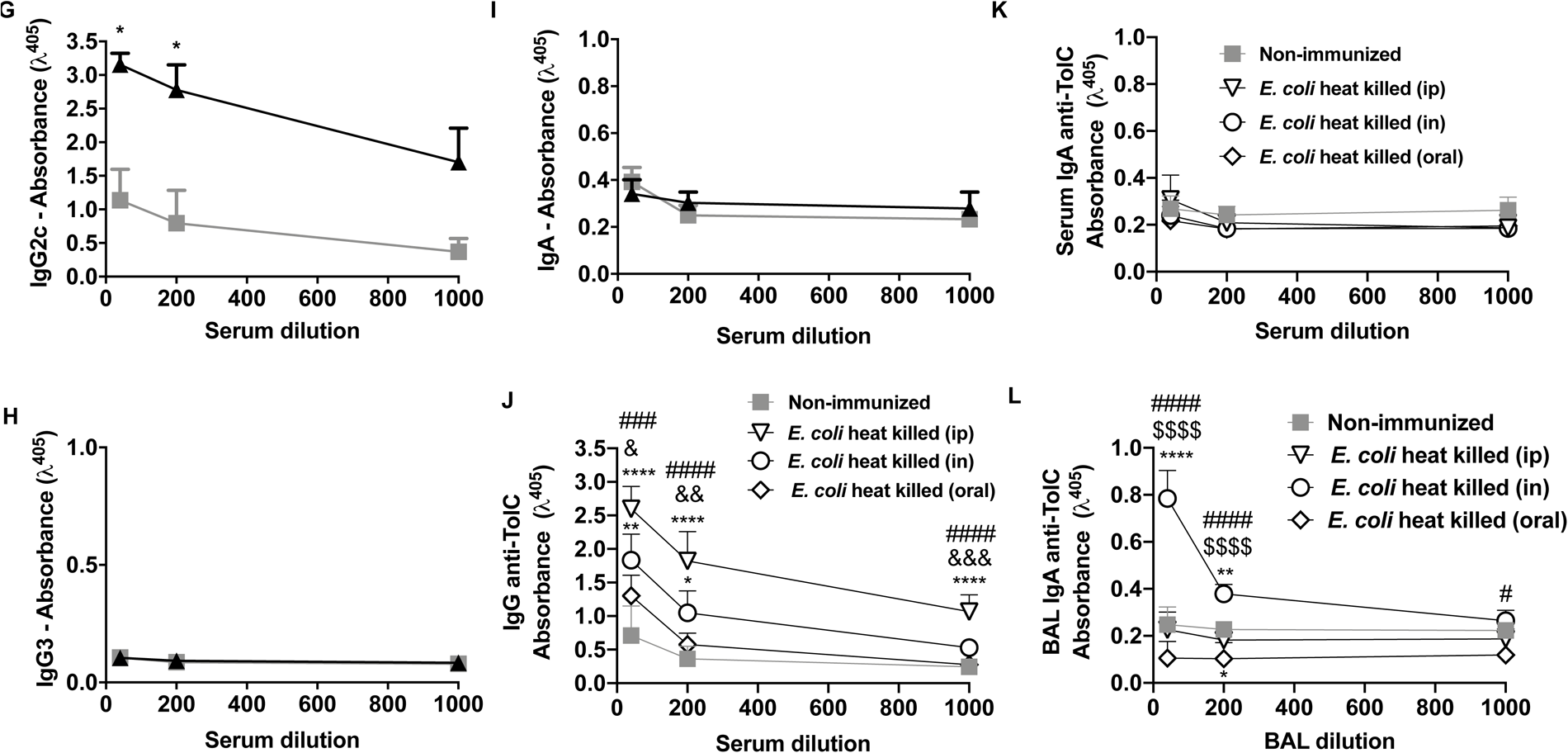
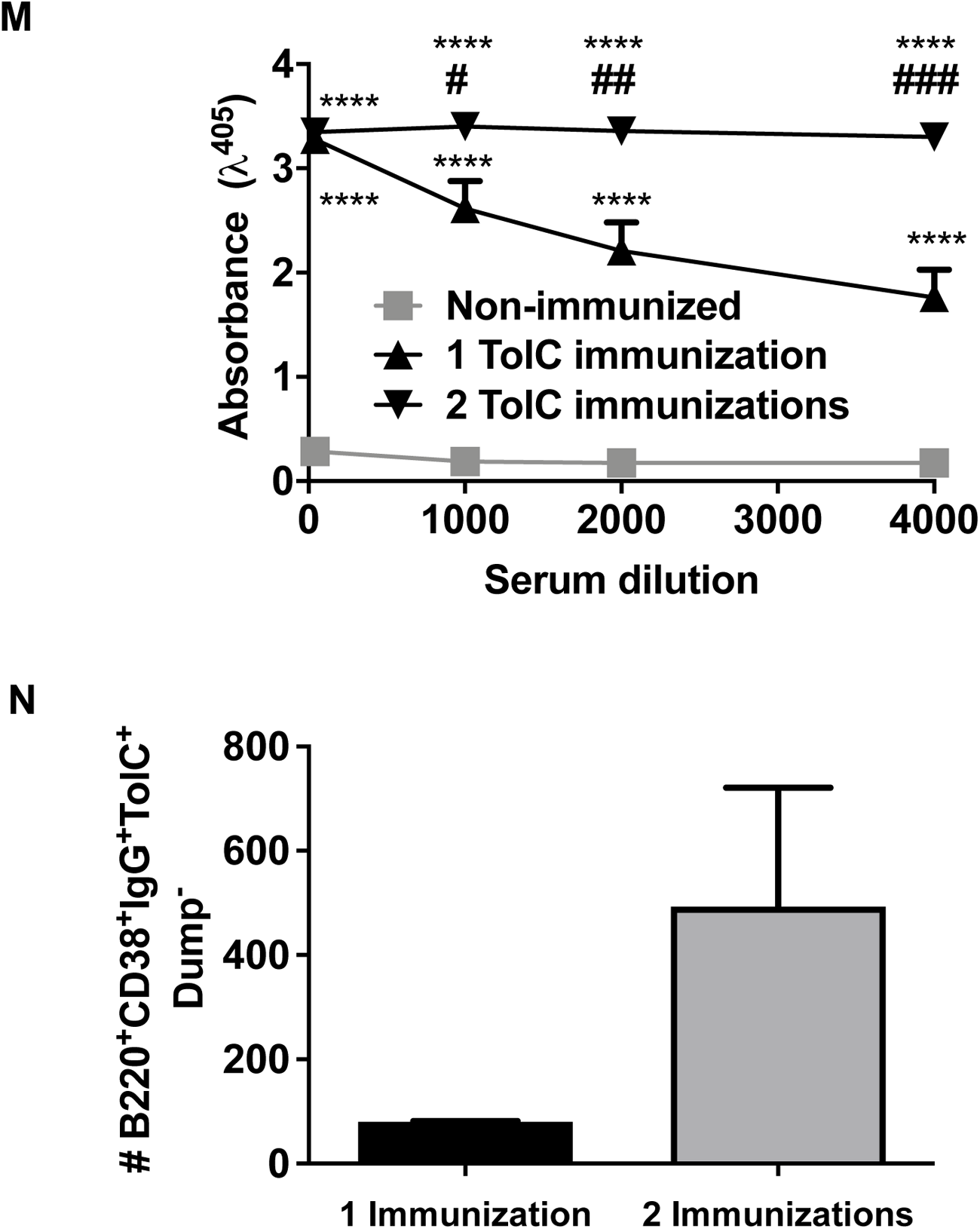
Immunization of mice with TolC recombinant protein induces B cell activation and antibody production. (A) C57/BL6 serum (1:120) IgM and (B) IgG anti-TolC from 3 to 35 days post TolC or no immunization. *** P <0.001 compared to non-immunized 7 days. **** P <0.0001 compared to the non-immunized one on the respective day. (C) C57/BL6 serum (1:40, 1:200 and 1:1000) anti-TolC IgM at 7 dpi and (D) anti-TolC IgG at 14 dpi compared to non-immunized and Germ-free mice. ** P <0.01 compared to non-immunized mouse serum at 1:40. **** P <0.0001 compared to “non-immunized”. C57/BL6 mice N=8-9. Germ-free mice N=2. (E) C57/BL6 anti-TolC serum IgG1, (F) IgG2b, (G) IgG2c, (H) IgG3 and (I) IgA 14days post TolC immunization. (J) C57/BL6 serum (1:40, 1:200 and 1:1000) anti-TolC IgG and (K) IgA at 14 days after the third immunization with heat killed whole cell *E. coli* (intranasal, intraperitoneal and oral). (L) C57/BL6 BAL (1:40, 1:200 and 1:1000) anti-TolC IgA at 14 days after the third immunization with heat killed whole cell *E. coli* (intranasal, intraperitoneal and oral). * P <0.05, ** P <0.01, **** P <0.0001 compared to non-immunized mouse serum at the respective sample dilution. ^&^P <0.05, ^&&^P <0.01, ^&&&^P <0.001 compared to the respective sample dilution of mouse immunized intranasally. #P <0.05, ^##^P <0.01, ^####^P <0.001 compared to the respective sample dilution of mouse immunized orally. ^$$$$^P <0.0001 compared to the respective sample dilution of mouse immunized intraperitoneally. C57/BL6 mice N=5-6. (M) BALB/c serum (1:40, 1:1000, 1:2000 and 1:4000) anti-TolC IgG at 14 days post 1 or 2 immunizations compared to non-immunized mice. (N) Number of TolC specific memory B cells in spleens and total lymph nodes of BALB/c mice immunized once or twice with TolC, evaluated by flow cytometry. **** P <0.0001 compared to the non-immunized one on the respective serum dilution. ^#^ P <0.05, ^##^ P <0.01, ^####^ P <0.0001 compared to mice immunized once with TolC. BALB/c mice N=3.

One or two immunizations of mice with whole cell heat-inactivated *E. coli* with a 14-day interval were not able to elicit anti-TolC IgG, independent of the administration route (intranasal, intraperitoneal or oral) (data not shown). However, when mice were immunized three times with heat-inactivated *E. coli* either intraperitoneally or intranasally, there was an increase in serum anti-TolC IgG titers, as compared to non-immunized mice (Figure 5J). Oral immunization with whole cell heat-inactivated *E. coli* did not lead to significant increase of serum anti-TolC IgG titers (Figure 5J). *E. coli* immunizations did not elicit serum anti-TolC IgA in any route tested (Figure 5K), but intranasal administration elicited BAL anti-TolC IgA (Figure 5 L).

BALB/c mice were also immunized with TolC protein once or twice, with a 14-day interval, and 14 days after the last challenge anti-TolC IgG titers and TolC specific memory B cells were assessed. A TolC booster was able to increase anti-TolC IgG production and numbers of TolC-specific memory B cells (Figure 5M and N, Figure S3 C and D).

Altogether, these results show TolC is immunogenic in mice and elicits memory B cells and specific IgA and IgG antibodies, with a predominance of subclasses IgG1 and IgG2.

### TolC-immune serum binds to *E. coli* and exerts protective effects

Serum IgG from mice immunized once or twice with TolC protein binds to live *E. coli* more than IgG from non-immunized mice (Figure 6A). We also tested whether anti-TolC IgG would bind to TolC recombinant protein as well bacteria cell extract proteins through Western blotting. We observed that TolC immune serum IgG binds to the TolC-GFP Tagged recombinant protein (81 kDa), as well as to *E. coli* TolC (54 kDa) and the homolog protein in *K. quasipneumoniae* (53 kDa), but not *P. aeruginosa* (53 kDa) or *A. baumannii* (50 kDa) (Figure 6 B).

**Figure 6:**
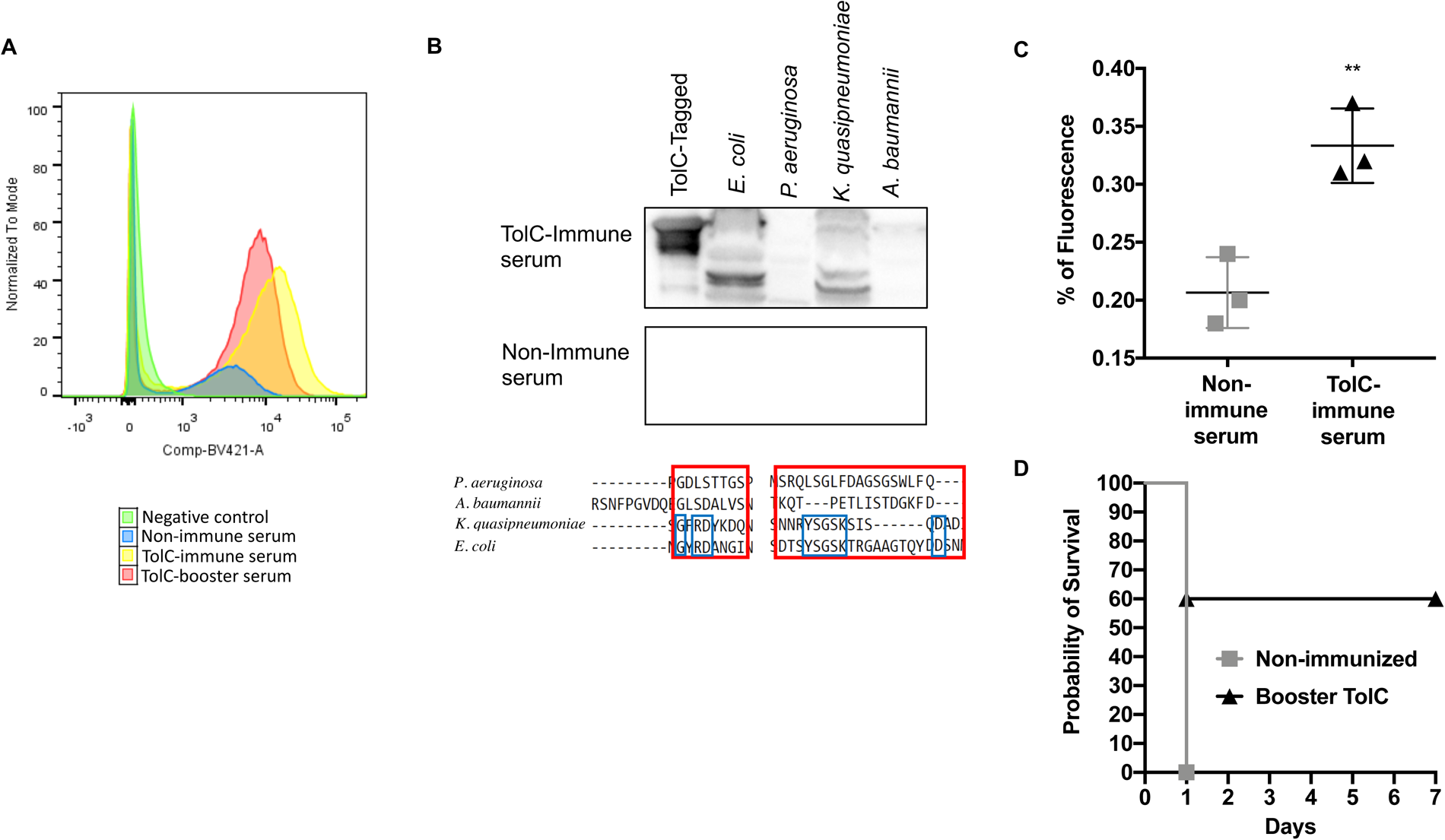
TolC-antibody bind to live *E. coli,* improves bacterial macrophage uptake and protects from *E. coli* infection. (A) IgG from serum of non-immune, or TolC immunized mice (once or booster) bound to live *E. coli* detected by flow cytometry. (B) IgG from serum of non-immune, or TolC immunized mice once bound to TolC-Tagged recombinant protein or protein extracts from *E. coli*, *P. aeruginosa*, *K. quasipneumoniae*, *A. baumannii*. Aligned sequences showing conserved residues (blue squares) in the extracellular regions (red squares) of TolC *E. coli* protein and homolog proteins from *P. aeruginosa*, *K. quasipneumoniae*, *A. baumannii.* (C) TolC immune serum incubated with live *E. coli* stained with Syto9 increases bacterial uptake by macrophage, detected by increased fluorescence in macrophages, through flow cytometry. ** P <0.01 compared to serum from non-immunized mice. BALB/c mice N=3. (D) Survival rate of mice either non-immunized or immunized and boosted with TolC infected with 2 x LD_50_ of *E. coli*. C57/BL6 male mice N=5-6.

Most importantly, TolC mouse immune serum increased *E. coli* macrophage uptake *in vitro* when compared to non-immune serum (Figure 6 C), and mice immunization with TolC was able to protect mice from *E. coli* infection by reducing mouse mortality in 60% (Figure 6 D).

## Discussion

Antimicrobial resistance has been declared a global threat by the WHO, given the rapid emergence and spread of antimicrobial resistant strains, accompanied by a lack of new antimicrobials under development ^4^. One of the mechanisms bacteria have developed that helped them overcome antimicrobials is the overexpression of efflux pumps, that leads to decreased cellular drug accumulation and is thought to contribute to multidrug resistance ^10,11^. *E. coli* is among the MDR bacterial pathogens considered by the WHO as a critical priority for drug research and development ^35^. Although *E. coli* is a commensal gut bacterium, some pathogenic serotypes are common causes of urinary tract infection, as well as bacteremia, neonatal meningitis and food-related outbreaks. The use of antimicrobials has been associated with an increase of MDR phenotypes overexpressing efflux pumps, especially AcrABTolC ^35–37^. Therefore, researching the immunogenicity of TolC, the outer membrane protein of this and other efflux pumps, may be of great interest.

An *in silico* study of outer membrane proteins from *Shigella flexneri* has demonstrated predicted B and T cell epitopes in TolC, considering this protein as a potential vaccine candidate^38^. Here we demonstrate that TolC from *E. coli* presents extracellular regions with predicted epitopes to human antibodies. Epitope prediction has been helping identify potential regions of interaction with molecules of the immune system. B-cell epitope prediction has direct implications in predicting the humoral response to the epitope. Most B-cell epitopes are discontinuous, composed of amino acid residues that reside in different regions of the protein and are brought together by protein folding. Ellipro, DiscoTope, and Epitopia, which are based on the 3D structure of a protein, identified the same epitope regions. The work of Borley *et al*. demonstrated that using three predictors and extracting the consensus reduces the false positive rate ^39^. The same methodology was applied for linear epitope prediction with BepiPred-2.0, ABCPred, and Epitopia from the TolC sequence. The predicted epitope regions were identified as antigenic, showing flexibility and propensity to interact with other proteins ^40^. The prediction results of interaction with class II MHC molecules for exposed epitopes indicated low interaction. However, 115 epitopes were predicted to have a high affinity for MHC II (IC_50_ <50 nM). These results suggest that antigen recognition by B cells and activation of helper T lymphocytes occur from different regions of the protein.

TolC and their homologs have emerged as interesting targets against Gram-negative bacteria ^17^ and it has been shown this protein is able to elicit antibodies in immunized rabbits ^18^. However, detailed study about the TolC ability to elicit innate and adaptive immune responses have not been addressed so far. Here we initially showed the presence of TolC antibodies in human plasma and found higher amounts of anti-TolC IgA and IgG than anti-TolC IgM, indicating *E. coli* TolC is an antigen recurrently presented to the human immune system. Accordingly, it has been shown that serum IgG antibodies are induced after infection or translocation of gut commensal *E. coli*, a bacteria with great capacity to translocate^41,42^. On ICU patients that presented recent Gram-negative infection there was a slight increase of anti-TolC IgM, when compared to control participants. This was not observed when comparing anti-LPS IgM or IgG, in between infected and control participants, suggesting TolC is more immunogenic than LPS. There was no difference in anti-TolC IgM or IgG plasma production regarding the participant gender or age. As far as we know, this is the first-time human antibodies against TolC have been identified.

To investigate how TolC is able to elicit antibodies, we applied *in vitro*, *ex vivo* and *in vivo* approaches. *In vitro*, macrophages recognize TolC recombinant protein, through mechanisms that are still unclear, and were stimulated to produce NO, IL-6 and TNF-α. This does not seem to be an effect of LPS contamination of the recombinant TolC protein produced in *E. coli*, since (i) the presence of LPS in 100 μg/mL of TolC recombinant protein was 10 times below the concentration permitted in injectable solutions ^43^, (ii) polymyxin did not interfere with NO production induced by TolC and (iii) mice immunization with this protein does not elicit LPS antibodies.

TolC immunization of mice led to increased lymph node T cell proliferation and IFN-γ secretion upon an *ex vivo* antigen recall. Previous studies have shown that immunization of rabbits with *E. coli* TolC recombinant protein or immunization with *Edwardsiella tarda* whole cell bacteria followed by TolC recombinant protein immunization of mice increased anti-TolC antibody production^17,18^. Here we demonstrate that one immunization of C57BL/6 mice with TolC emulsified in alum does not affect serum anti-TolC IgA production, mouse weight or survival, but leads to increased anti-TolC IgM at 7 dpi, and anti-TolC IgG at 14 dpi. IgG anti-TolC is maintained at high levels until at least 35 dpi. BALB/c mouse immunization with TolC also leads to increased specific IgG serum production, and the booster improves both anti-TolC IgG and specific memory B cell amounts.

IFNγ has been associated with B cell IgG2 secretion in mice and humans ^44,45^. Here we observed that TolC immunization and re-stimulation *in vitro* induced increased IFNγ production by lymph node cells and higher titers of IgG1, IgG2b and IgG2c, but not IgG3. Accordingly, human anti-TolC IgG1 and IgG2 were more prevalent than IgG3 and IgG4 in plasma, suggesting IFNγ induced by TolC can be involved in shaping this response. IgG1 represents a bigger percentage of total IgG in adult serum, has bigger biological half-life and, as well as IgG3, is generally produced in response to proteins ^46^, nonetheless anti-TolC IgG2 seemed more prevalent in human and mice serum. In fact, IgG1 and IgG3 are thought to be potent triggers of effector mechanisms, and IgG2 can activate complement at high densities of surface antigens, as is the case for polysaccharides and the outer membrane protein TolC ^46,47^. In a previous study, rabbit antibodies against TolC were able to bind to *E. coli* TolC extracted proteins and reduce bacteria cell growth *in vitro*, when used in the presence of 1/8 of chloramphenicol MIC^18^. Another study has shown that serum from mice immunized with *A. baumannii* TolC recombinant protein increased bacterial complement-dependent killing and opsonophagocytosis ^48^. Accordingly, we observed that anti-TolC IgG from mouse serum bound to TolC from *E. coli* protein extracts as well as to live *E. coli.* IgG from *E. coli* TolC recombinant protein immunized mice also bound to protein extracts from the pathogenic bacteria *K. quasipneumoniae*, probably due to higher similarity of their TolC extracellular residues, but not to *P. aeruginosa* or *A. baumannii*. Noteworthy, we show that mouse serum antibodies against the *E. coli* TolC, increased *E. coli* uptake by macrophage and also increased mice survival after *E. coli* lethal infection by 60%. As far as we know, this is the first time that it is shown that anti-TolC antibodies have a protective role against *E. coli* infection *in vivo*.

It was interesting to note that lymph node T cells from naïve mice stimulated *in vitro* with TolC also showed an increase in IFN-γ production, at least in the highest concentration of TolC. Non-immunized mice also presented anti-TolC IgM and IgG antibodies, when serum was diluted at 1:40, which was not seen in Germ-free mice. These findings suggest that mice microbiota could have a role in TolC elicited antibodies, probably due to the *E. coli* translocation ability ^41,42^. The presence of anti-TolC IgM, IgG and IgA in healthy human subjects corroborates this hypothesis. To test if *E. coli* introduction through different routes could elicit anti-TolC antibodies, naïve mice were immunized with heat killed bacteria via oral, intraperitoneal or intranasal routes. Only after the third intraperitoneal or intranasal immunization with *E. coli* there was a significant increase in serum anti-TolC IgG, suggesting *E. coli* introduction through different routes has different potential to induce antibody production against TolC, with higher tolerance in oral administration, as expected. Although humans presented high titers of anti-TolC IgA in the serum, immunization of mice with TolC recombinant protein or heat killed *E. coli* in all three routes tested did not lead to increase in serum anti-TolC IgA. Nonetheless, intranasal bacterial challenge led to increased levels of anti-TolC IgA in the lungs. The fact that humans present mostly monomeric serum IgA, as opposed to mice that present mostly polymeric IgA ^49^, could help explain the difference in between human and mice serum anti-TolC IgA levels.

In this study, we report novel observations regarding the immunogenic role of TolC, an *E. coli* outer membrane protein that composes efflux pumps participate in bacterial multidrug resistance. We identified the extracellular regions of TolC that are more likely to be epitopes for human antibodies and demonstrated that human plasma presents IgM, IgG and IgA against TolC, with a higher prevalence of IgG2 amongst IgG subclasses. There was also a slight increase in human anti-TolC IgM amounts following infection with Gram-negative bacteria. To understand how TolC stimulates the immune system leading to antibody production, we assessed *in vitro* and *in vivo* mouse models. TolC is recognized by macrophages stimulating them to secrete proinflammatory mediators important for initiating the immune response. An *ex vivo* TolC re-stimulation led to T cell proliferation and IFN-γ production. Mice immunized with TolC recombinant protein, or with intranasal heat killed *E. coli*, produced high amounts of anti-TolC IgG, with a prevalence of IgG1 and IgG2. IgG anti-TolC was able to bind live *E. coli* and to TolC protein from *E. coli* and *K. quasipneumoniae* cell extracts. More importantly, murine TolC-immune serum increased *E. coli* macrophage uptake and protected mice from a lethal infection. Altogether, our findings indicate that TolC is an immunogenic protein leading to antibody production in humans and mice, and these antibodies might participate in the immune response against *E. coli*. Further experiments are needed to evaluate the activity of anti-TolC antibodies against antimicrobial resistant Gram-negative bacteria.

## Acknowledgments

We thank Dr Leda Quercia Vieira and Dr Flaviano dos Santos Martins from Instituto de Ciências Biológicas of the Universidade Federal de Minas Gerais (UFMG) for kindly donating serum from the Germ-free mice from their Animal Facility. This study was supported by grants from Pew Charitable Trusts Funds, Brazilian Council for Scientific and Technological Development (CNPq) and Fundação Carlos Chagas de Amparo à Pesquisa do Estado do Rio de Janeiro (FAPERJ). OST, CSLV and BACS were supported by fellowships from CAPES; ARSC, RBF, BLS, PSAJ were supported by fellowships from CNPq; CNL, FJG, SSL and BACS were supported by fellowships from FAPERJ.

## Notes

### Competing Interest Statement

The authors have declared no competing interest.

