## Supplementary figures and images for "The *Escherichia coli* TolC efflux pump protein is immunogenic and elicits protective antibodies"

### Supplemental figures

#### Slide 1
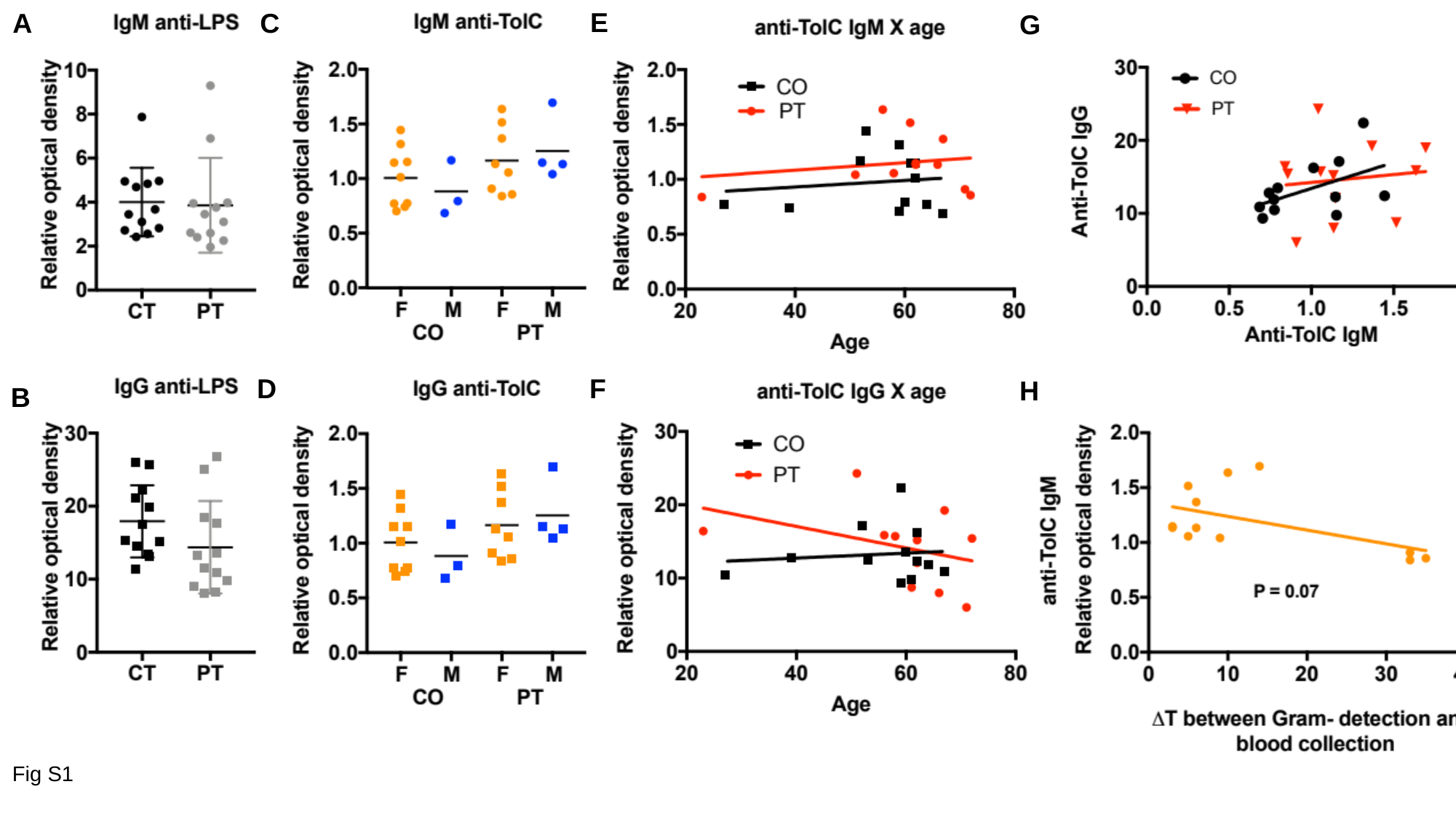

E
A
C
G
D
F
H
B
Fig S1

#### Slide 2
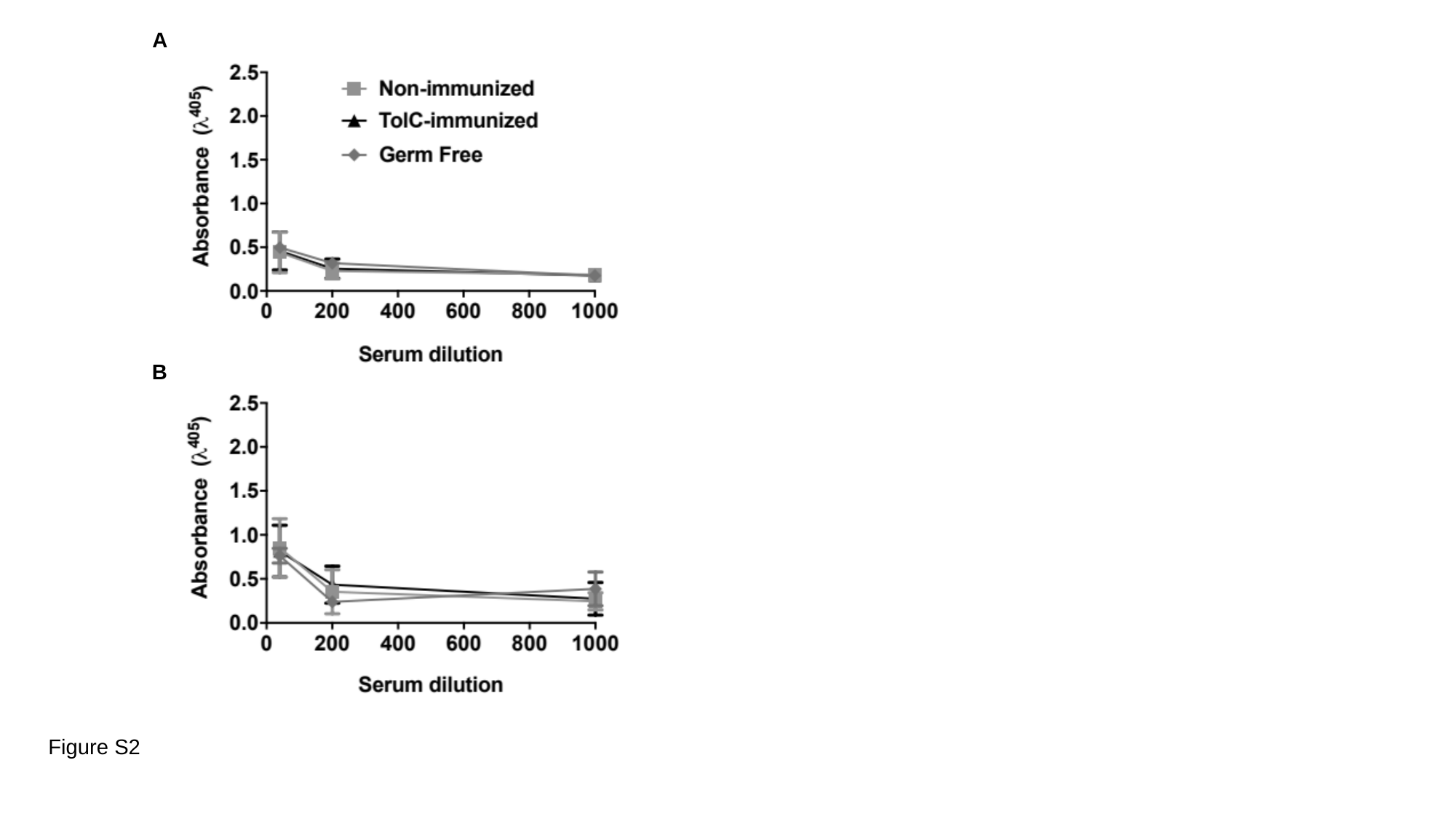

A
B
Figure S2

#### Slide 3
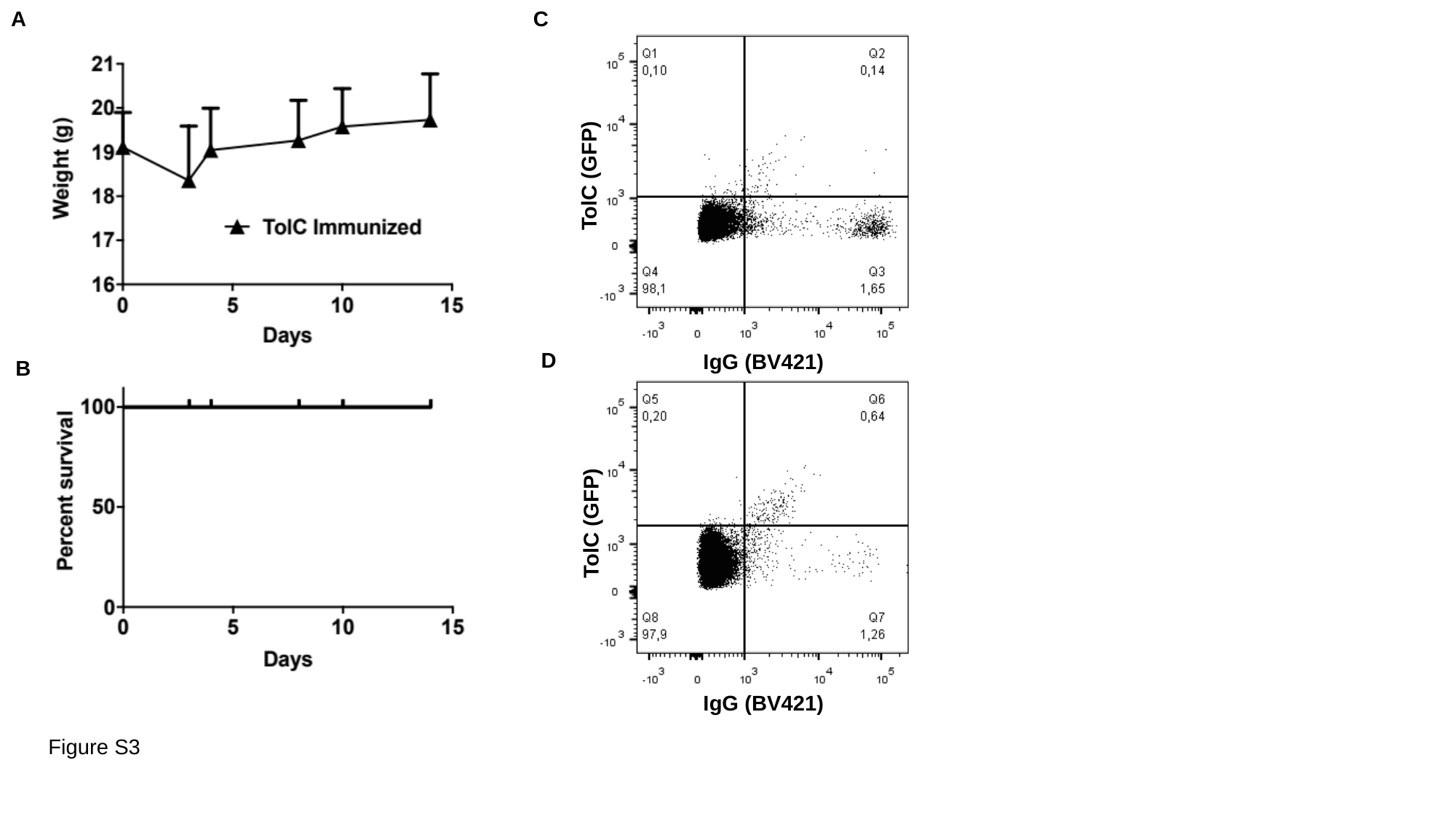

A
C
TolC (GFP)
D
IgG (BV421)
B
TolC (GFP)
IgG (BV421)
Figure S3
