## Supplemental table for "The *Escherichia coli* TolC efflux pump protein is immunogenic and elicits protective antibodies"

**Supplementary Table S1.** T Cell Epitope Prediction (MHC II).

| TolC *E. coli* | | | | | |
|---|---|---|---|---|---|
| **IC50** | **Allele** | **start** | **end** | **core_peptide** | **Peptide** |
| 3.40 | HLA-DRB3*02:02 | 124 | 138 | LILNTATAY | QQTLILNTATAYFNV |
| 4.10 | HLA-DRB3*02:02 | 123 | 137 | LILNTATAY | DQQTLILNTATAYFN |
| 4.60 | HLA-DRB3*02:02 | 125 | 139 | LILNTATAY | QTLILNTATAYFNVL |
| 5.40 | HLA-DRB1*07:01 | 264 | 278 | LDLTASTGI | DGHLPTLDLTASTGI |
| 5.40 | HLA-DRB1*07:01 | 265 | 279 | LDLTASTGI | GHLPTLDLTASTGIS |
| 5.50 | HLA-DRB3*02:02 | 122 | 136 | LILNTATAY | TDQQTLILNTATAYF |
| 5.90 | HLA-DRB1*07:01 | 266 | 280 | LDLTASTGI | HLPTLDLTASTGISD |
| 6.80 | HLA-DRB1*07:01 | 267 | 281 | LDLTASTGI | LPTLDLTASTGISDT |
| 7.40 | HLA-DRB1*07:01 | 268 | 282 | LDLTASTGI | PTLDLTASTGISDTS |
| 7.90 | HLA-DRB1*07:01 | 269 | 283 | LDLTASTGI | TLDLTASTGISDTSY |
| 8.30 | HLA-DRB3*02:02 | 432 | 446 | LALNNALSK | QDLLALNNALSKPVS |
| 8.50 | HLA-DRB3*01:01 | 116 | 130 | YQTDQQTLI | QDVTYQTDQQTLILN |
| 8.60 | HLA-DRB3*01:01 | 117 | 131 | YQTDQQTLI | DVTYQTDQQTLILNT |
| 9.00 | HLA-DRB3*01:01 | 115 | 129 | YQTDQQTLI | IQDVTYQTDQQTLIL |
| 9.30 | HLA-DRB3*02:02 | 433 | 447 | LALNNALSK | DLLALNNALSKPVST |
| 9.80 | HLA-DRB3*01:01 | 118 | 132 | YQTDQQTLI | VTYQTDQQTLILNTA |
| 10.60 | HLA-DRB5*01:01 | 240 | 254 | LSLLQARLS | KRNLSLLQARLSQDL |
| 11.40 | HLA-DRB3*02:02 | 431 | 445 | LALNNALSK | EQDLLALNNALSKPV |
| 11.50 | HLA-DRB5*01:01 | 238 | 252 | LSLLQARLS | AEKRNLSLLQARLSQ |
| 11.50 | HLA-DRB5*01:01 | 239 | 253 | LSLLQARLS | EKRNLSLLQARLSQD |
| 11.70 | HLA-DRB3*02:02 | 351 | 365 | NNINASISS | SSFNNINASISSINA |
| 13.00 | HLA-DRB1*07:01 | 178 | 192 | YDTVLANEV | NARAQYDTVLANEVT |
| 13.40 | HLA-DRB3*02:02 | 121 | 135 | LILNTATAY | QTDQQTLILNTATAY |
| 13.60 | HLA-DRB1*07:01 | 177 | 191 | YDTVLANEV | QNARAQYDTVLANEV |
| 14.00 | HLA-DRB3*02:02 | 350 | 364 | NNINASISS | RSSFNNINASISSIN |
| 14.10 | HLA-DRB1*07:01 | 270 | 284 | LDLTASTGI | LDLTASTGISDTSYS |
| 14.40 | HLA-DRB3*02:02 | 126 | 140 | LILNTATAY | TLILNTATAYFNVLN |
| 14.60 | HLA-DRB3*02:02 | 352 | 366 | NNINASISS | SFNNINASISSINAY |
| 14.80 | HLA-DRB1*07:01 | 179 | 193 | YDTVLANEV | ARAQYDTVLANEVTA |
| 15.00 | HLA-DRB1*07:01 | 300 | 314 | VGLSFSLPI | NMGQNKVGLSFSLPI |
| 16.70 | HLA-DRB5*01:01 | 241 | 255 | LSLLQARLS | RNLSLLQARLSQDLA |
| 17.00 | HLA-DRB3*02:02 | 349 | 363 | NNINASISS | VRSSFNNINASISSI |
| 17.20 | HLA-DRB3*01:01 | 114 | 128 | YQTDQQTLI | GIQDVTYQTDQQTLI |
| 17.40 | HLA-DRB1*07:01 | 301 | 315 | VGLSFSLPI | MGQNKVGLSFSLPIY |
| 17.50 | HLA-DRB3*02:02 | 430 | 444 | LALNNALSK | NEQDLLALNNALSKP |
| 18.10 | HLA-DRB1*07:01 | 180 | 194 | YDTVLANEV | RAQYDTVLANEVTAR |
| 18.40 | HLA-DRB3*02:02 | 412 | 426 | YLINQLNIK | RYNYLINQLNIKSAL |
| 18.80 | HLA-DRB3*01:01 | 119 | 133 | YQTDQQTLI | TYQTDQQTLILNTAT |
| 19.20 | HLA-DRB5*01:01 | 237 | 251 | LSLLQARLS | EAEKRNLSLLQARLS |
| 19.40 | HLA-DRB1*07:01 | 302 | 316 | VGLSFSLPI | GQNKVGLSFSLPIYQ |
| 21.10 | HLA-DRB3*02:02 | 353 | 367 | INASISSIN | FNNINASISSINAYK |
| 21.30 | HLA-DRB3*02:02 | 434 | 448 | LALNNALSK | LLALNNALSKPVSTN |
| 21.80 | HLA-DRB5*01:01 | 356 | 370 | SISSINAYK | INASISSINAYKQAV |
| 21.90 | HLA-DRB3*02:02 | 411 | 425 | YLINQLNIK | ARYNYLINQLNIKSA |
| 22.70 | HLA-DRB4*01:01 | 412 | 426 | YLINQLNIK | RYNYLINQLNIKSAL |
| 22.90 | HLA-DRB5*01:01 | 357 | 371 | ISSINAYKQ | NASISSINAYKQAVV |
| 25.30 | HLA-DRB1*15:01 | 240 | 254 | LSLLQARLS | KRNLSLLQARLSQDL |
| 25.40 | HLA-DRB3*02:02 | 413 | 427 | YLINQLNIK | YNYLINQLNIKSALG |

| | | | | | |
|---|---|---|---|---|---|
| 25.50 | HLA-DRB1*15:01 | 358 | 372 | INAYKQAVV | ASISSINAYKQAVVS |
| 26.50 | HLA-DRB1*07:01 | 181 | 195 | YDTVLANEV | AQYDTVLANEVTARN |
| 26.60 | HLA-DRB4*01:01 | 413 | 427 | YLINQLNIK | YNYLINQLNIKSALG |
| 26.70 | HLA-DRB1*07:01 | 303 | 317 | VGLSFSLPI | QNKVGLSFSLPIYQG |
| 26.90 | HLA-DRB4*01:01 | 411 | 425 | YLINQLNIK | ARYNYLINQLNIKSA |
| 27.60 | HLA-DRB1*15:01 | 239 | 253 | LSLLQARLS | EKRNLSLLQARLSQD |
| 28.00 | HLA-DRB1*15:01 | 302 | 316 | VGLSFSLPI | GQNKVGLSFSLPIYQ |
| 28.60 | HLA-DRB1*15:01 | 238 | 252 | LSLLQARLS | AEKRNLSLLQARLSQ |
| 28.70 | HLA-DRB1*07:01 | 131 | 145 | FNVLNAIDV | TATAYFNVLNAIDVL |
| 28.90 | HLA-DRB5*01:01 | 99 | 113 | WRALTLQEK | DMSKWRALTLQEKAA |
| 29.10 | HLA-DRB5*01:01 | 100 | 114 | WRALTLQEK | MSKWRALTLQEKAAG |
| 29.20 | HLA-DRB3*02:02 | 410 | 424 | YLINQLNIK | NARYNYLINQLNIKS |
| 30.50 | HLA-DRB5*01:01 | 355 | 369 | SISSINAYK | NINASISSINAYKQA |
| 30.60 | HLA-DRB1*15:01 | 357 | 371 | ISSINAYKQ | NASISSINAYKQAVV |
| 30.60 | HLA-DRB1*07:01 | 132 | 146 | FNVLNAIDV | ATAYFNVLNAIDVLS |
| 30.70 | HLA-DRB4*01:01 | 414 | 428 | YLINQLNIK | NYLINQLNIKSALGT |
| 30.90 | HLA-DRB1*07:01 | 94 | 108 | FDMSKWRAL | TQSIFDMSKWRALTL |
| 30.90 | HLA-DRB5*01:01 | 411 | 425 | YNYLINQLN | ARYNYLINQLNIKSA |
| 31.20 | HLA-DRB1*15:01 | 22 | 36 | MQVYQQARL | AENLMQVYQQARLSN |
| 31.40 | HLA-DRB1*15:01 | 303 | 317 | VGLSFSLPI | QNKVGLSFSLPIYQG |
| 31.40 | HLA-DRB5*01:01 | 410 | 424 | YNYLINQLN | NARYNYLINQLNIKS |
| 31.60 | HLA-DRB3*01:01 | 220 | 234 | FKTDKPQPV | VENFKTDKPQPVNAL |
| 31.60 | HLA-DRB1*07:01 | 133 | 147 | FNVLNAIDV | TAYFNVLNAIDVLSY |
| 32.30 | HLA-DRB4*01:01 | 367 | 381 | VVSAQSSLD | KQAVVSAQSSLDAME |
| 32.30 | HLA-DRB5*01:01 | 242 | 256 | LSLLQARLS | NLSLLQARLSQDLAR |
| 32.50 | HLA-DRB4*01:01 | 368 | 382 | VVSAQSSLD | QAVVSAQSSLDAMEA |
| 32.70 | HLA-DRB1*15:01 | 359 | 373 | INAYKQAVV | SISSINAYKQAVVSA |
| 32.90 | HLA-DRB1*15:01 | 23 | 37 | MQVYQQARL | ENLMQVYQQARLSNP |
| 33.30 | HLA-DRB3*02:02 | 348 | 362 | NNINASISS | TVRSSFNNINASISS |
| 33.50 | HLA-DRB1*07:01 | 130 | 144 | FNVLNAIDV | NTATAYFNVLNAIDV |
| 33.70 | HLA-DRB5*01:01 | 358 | 372 | ISSINAYKQ | ASISSINAYKQAVVS |
| 34.00 | HLA-DRB1*03:01 | 183 | 197 | VLANEVTAR | YDTVLANEVTARNNL |
| 34.30 | HLA-DRB1*07:01 | 134 | 148 | FNVLNAIDV | AYFNVLNAIDVLSYT |
| 34.80 | HLA-DRB5*01:01 | 412 | 426 | YNYLINQLN | RYNYLINQLNIKSAL |
| 35.90 | HLA-DRB1*07:01 | 304 | 318 | VGLSFSLPI | NKVGLSFSLPIYQGG |
| 37.30 | HLA-DRB3*01:01 | 219 | 233 | FKTDKPQPV | NVENFKTDKPQPVNA |
| 37.60 | HLA-DRB1*07:01 | 135 | 149 | FNVLNAIDV | YFNVLNAIDVLSYTQ |
| 37.90 | HLA-DRB1*15:01 | 21 | 35 | MQVYQQARL | QAENLMQVYQQARLS |
| 38.00 | HLA-DRB1*07:01 | 95 | 109 | FDMSKWRAL | QSIFDMSKWRALTLQ |
| 38.20 | HLA-DRB5*01:01 | 98 | 112 | WRALTLQEK | FDMSKWRALTLQEKA |
| 38.50 | HLA-DRB1*07:01 | 182 | 196 | YDTVLANEV | QYDTVLANEVTARNN |
| 38.50 | HLA-DRB5*01:01 | 101 | 115 | WRALTLQEK | SKWRALTLQEKAAGI |
| 38.60 | HLA-DRB1*15:01 | 241 | 255 | LSLLQARLS | RNLSLLQARLSQDLA |
| 39.10 | HLA-DRB4*01:01 | 240 | 254 | NLSLLQARL | KRNLSLLQARLSQDL |
| 39.10 | HLA-DRB3*02:02 | 220 | 234 | FKTDKPQPV | VENFKTDKPQPVNAL |
| 39.20 | HLA-DRB1*07:01 | 47 | 61 | INEARSPLL | DAAFEKINEARSPLL |
| 39.40 | HLA-DRB1*03:01 | 182 | 196 | VLANEVTAR | QYDTVLANEVTARNN |
| 39.70 | HLA-DRB4*01:01 | 241 | 255 | NLSLLQARL | RNLSLLQARLSQDLA |
| 40.00 | HLA-DRB3*01:01 | 218 | 232 | FKTDKPQPV | LNVENFKTDKPQPVN |
| 40.20 | HLA-DRB4*01:01 | 410 | 424 | YLINQLNIK | NARYNYLINQLNIKS |
| 40.20 | HLA-DRB1*15:01 | 24 | 38 | MQVYQQARL | NLMQVYQQARLSNPE |
| 40.50 | HLA-DRB1*15:01 | 301 | 315 | VGLSFSLPI | MGQNKVGLSFSLPIY |

| | | | | | |
|---|---|---|---|---|---|
| 40.90 | HLA-DRB3*02:02 | 161 | 175 | QRFNVGLVA | QTTQRFNVGLVAITD |
| 41.40 | HLA-DRB1*03:01 | 184 | 198 | VLANEVTAR | DTVLANEVTARNNLD |
| 42.70 | HLA-DRB1*15:01 | 304 | 318 | VGLSFSLPI | NKVGLSFSLPIYQGG |
| 43.20 | HLA-DRB1*15:01 | 360 | 374 | INAYKQAVV | ISSINAYKQAVVSAQ |
| 43.30 | HLA-DRB4*01:01 | 366 | 380 | VVSAQSSLD | YKQAVVSAQSSLDAM |
| 44.00 | HLA-DRB3*02:02 | 429 | 443 | LALNNALSK | LNEQDLLALNNALSK |
| 45.10 | HLA-DRB1*07:01 | 96 | 110 | FDMSKWRAL | SIFDMSKWRALTLQE |
| 45.80 | HLA-DRB4*01:01 | 239 | 253 | NLSLLQARL | EKRNLSLLQARLSQD |
| 46.20 | HLA-DRB5*01:01 | 409 | 423 | YNYLINQLN | ANARYNYLINQLNIK |
| 47.70 | HLA-DRB3*01:01 | 221 | 235 | FKTDKPQPV | ENFKTDKPQPVNALL |
| 47.70 | HLA-DRB1*15:01 | 361 | 375 | INAYKQAVV | SSINAYKQAVVSAQS |
| 47.80 | HLA-DRB1*07:01 | 48 | 62 | INEARSPLL | AAFEKINEARSPLLP |
| 47.90 | HLA-DRB3*02:02 | 162 | 176 | QRFNVGLVA | TTQRFNVGLVAITDV |
| 49.90 | HLA-DRB4*01:01 | 238 | 252 | NLSLLQARL | AEKRNLSLLQARLSQ |
| 50.90 | HLA-DRB3*02:02 | 409 | 423 | YLINQLNIK | ANARYNYLINQLNIK |
